# CONVEX: Consensus Variant Pipeline for Exome Analysis

**DOI:** 10.1101/2023.10.07.561328

**Authors:** Ranjana Mariyam Raju, Ujjwal Prathap Singh, Prashanth Suravajhala

## Abstract

Benchmarking whole exome pipelines is crucial for evaluating and comparing their performance in variant calling and clinical significance annotation. It enables researchers and clinicians to assess the accuracy, sensitivity, and specificity of different pipelines and identify the most effective and reliable ones. In this study, we evaluated and compared the performance of our in-house consensus exome pipeline with a widely recognized gold standard Genome Analysis Toolkit (GATK) pipeline. Four datasets were used for evaluation, three 1000 Genome Project (1KGP) datasets and one Prostate cancer (PCa) Sample. The consensus pipeline consistently demonstrated a higher average transition-to-transversion (Ti/Tv) ratio, indicating enhanced precision in identifying single nucleotide variant (SNV) calls. This suggests that the consensus pipeline excels in effectively discerning true genetic variations from sequencing artefacts, particularly in the context of exome sequencing. Additionally, the pipeline exhibited increased sensitivity in detecting pathogenic and likely pathogenic variants in the PCa sample, further highlighting its efficacy in identifying clinically relevant variants. We also conducted a trio exome analysis with the use of two trio pipelines, *viz*. VarScan Trio and GATK joint calling pipelines on our erstwhile Congenital Pouch Colon (CPC) samples from our rare disease cohort which we published earlier and found that the GATK predicted a significantly higher number of variants compared to VarScan. From our pipeline, *viz*. CONVEX: Consensus Variant Pipeline for Exome Analysis we developed, our study demonstrates a large potential for trio-variant calling analysis paving the way for precision medicine. We discuss the impending benchmark results using the CONVEX.

## Introduction

Genomic analysis refers to the process of examining, interpreting, and understanding the information encoded in an organism’s DNA sequence. This can involve various techniques and approaches, such as identifying genes, detecting genetic variations and studying gene expression patterns. High-throughput genomic analysis has been made possible by the development of next-generation sequencing (NGS) technologies, which generate massive amounts of genomic data[1]. The main feature distinguishing NGS from Sanger sequencing is its ability to perform massively parallel sequencing of DNA or RNA fragments, allowing for the simultaneous analysis of multiple samples or targets[2].

Whole exome sequencing (WES) has several advantages over whole genome sequencing (WGS) in certain contexts, for example, cost and resource efficiency as it focuses on the protein-coding regions (exons) of the genome, which constitute only about 1-2% of the entire genome. WES specifically targets the exons, where most known disease-causing mutations occur, allowing researchers to focus on the most functionally relevant parts of the genome, increasing the likelihood of identifying causative mutations[3,4]. While the targeted approach can be particularly useful for diagnosing rare genetic disorders caused by mutations in these regions, the WES or targeted approaches generate a smaller dataset compared to WGS, making data analysis and storage more feasible, and particularly beneficial for clinical research laboratories with limited computational resources. While WES can also provide better sensitivity for detecting single nucleotide variants (SNVs) and small insertions/deletions (indels) in the coding regions compared to WGS, it has limitations, such as the inability to detect non-coding and structural variants, which may be relevant in certain diseases or research contexts[5]. As sequencing costs continue to decrease and bioinformatics tools improve, the use of WGS may become more widespread, offering a comprehensive view of the genome. In some cases, WGS may be more suitable for diagnosing rare diseases, as it can provide a more comprehensive view of the genome and identify pathogenic variants in non-coding regions.

On the other hand, whole exome-trio sequencing is a powerful tool in clinical genetic diagnosis and screening for hereditary disorders, particularly for patients with undiagnosed or rare disorders. This approach involves sequencing the exomes (the protein-coding regions of the genome) of three related individuals: a proband (an individual with the disease or trait of interest), and their unaffected parents. By comparing the exomes of the trio, researchers can identify rare variants that are inherited from one or both parents which may be contributing to the proband’s disease. While this method relies on high-quality genome-sequencing data and sophisticated data-interpretation approaches, the results are returned to the physician in a concise report with an in-depth phenotype-driven interpretation. The patients have the option to decide whether the report should include secondary and incidental findings. The integration of DNA, RNA, and protein data may refine the therapeutic stratification of individual patients, potentially increasing the success rate of precision cancer therapy[6]. For example, in the context of epilepsy, obtaining a genetic diagnosis could help families in terms of informing further reproductive decisions, providing answers and preventing further costly investigations[7]. Apart from the aforementioned utilities of WES, the advantages of using whole exome-trio analysis for rare diseases include (a) identification of *de novo* mutations, which allows the identification of *de novo* mutations (DNMs) that are not present in the parents but have arisen in the affected individual[8]. (b) Increased diagnostic yield and enhanced interpretation wherein the case-parent trio design enables better interpretation of the genetic variants identified, as it allows for comparison of affected individual’s genetic data with that of their parents[9]. This helps in distinguishing between inherited and *de novo* mutations, and in determining the potential pathogenicity of the identified variants. (c) Integration with other data types, wherein this could be combined with other data types, such as RNA sequencing and proteomics, to provide a more comprehensive understanding of the molecular basis of rare diseases and guide targeted treatment in precision medicine[6].

Benchmarking is a process of comparing the performance of a system or process against a standard or best practice. Benchmarking in whole exome-trio sequencing pipelines can be performed by comparing the results obtained from the analyses to a known set of high-quality reference data allowing the identification of errors or biases in the pipeline thereby improving its performance. One way to perform benchmarking is to use a uniform tumour-normal sample pair and compare sequencing pipelines at different centres using a standardised algorithm[10]. This can help identify the impact of varying library preparation methods and sequencing coverage metrics on downstream results. Another way to perform benchmarking is to assess the performance of varying prioritisation tools for germline causal variants from WES data[11]. In this case, the diagnostic yield of each tool can be evaluated based on the first, fifth and tenth rankings of the causal variant. Furthermore, it helps evaluate the impact of changes or updates in the pipeline, enabling the researchers to produce effective results that are of high quality which can then be used for downstream analysis and interpretation. In this work, we performed a trio exome analysis with the use of two trio pipelines, *viz*. VarScan Trio and GATK joint calling pipelines on Congenital Pouch Colon (CPC) samples from our rare disease cohort([12],[13],[14] and Prostate cancer (PCa)[15] which we have published earlier. We attempted to compare the benchmarking results by analysing four datasets with a high confidence variant call set by the Genome in a Bottle (GIAB) consortium besides considering a truth set for pipeline performance validation to confirm their association with disease-causing mutations in CPC and PCa. We further discuss the impending challenges in predicting pathogenic variants of significance from our benchmarking results on these datasets.

## Materials and Methods

### Benchmarking datasets for consensus pipeline

Four datasets were analysed with one dataset having a high confidence variant call set. FASTQ files of human exome HapMap/1000 Genomes project (1KGP) CEU female NA12878 (accession No.: SRR098401) were downloaded from NCBI-Sequence Read Archive (SRA-https://www.ncbi.nlm.nih.gov/sra last accessed on June 10, 2023). The WES of NA12878 was performed using the HiSeq2500 platform and Nextera Rapid Capture Exome and Expanded Exome kit. The human reference genomes GRCh38 were downloaded from the Ensembl[16]. Next, a NA12878 high confidence call set to version 4.2.2 by the GIAB consortium was used for pipeline performance validation. The variant set along with a BED file was downloaded from NCBI. This GIAB variant set, created by integrating 14 different datasets from five sequencers, is a ‘gold standard’ variant dataset publicly available for systematic comparison of variant callers. For the analysis of pathogenic variant distribution, pathogenic, likely pathogenic variants and variants of uncertain/unknown significance (VoUS) with non-conflicting interpretations were selected from ClinVar v.20230520. For added validation, two genomes from the 1KGP-CHS (East Asian Ancestry) and FIN (European Ancestry) populations were taken (SRA IDs: ERR032022, ERR031490) apart from the PCa samples from the cohort used in our Systems Genomics lab as discussed earlier.

### Trio datasets

For the trio-exome analysis, we employed the VarScan Trio and GATK Joint Variant Calling Pipeline, one GIAB trio of NA12878 proband with father NA12891 and mother NA12892. Father and Mother FASTQ files were downloaded from 1KGP (A map of human genome variation from population-scale sequencing 2010; SRA IDs: SRR098359, ERR034529; Table 1). Furthermore, two of the original CPC samples (Z12 and Z19 family) from a previous study conducted in our Systems Genomics lab were taken for conducting a retrospective study on the rare variants found using two trio variant calling pipelines with updated ClinVar records.

**Table 1:** Datasets used for analyses.

| Benchmarking datasets |  |  |  |
| --- | --- | --- | --- |
| Dataset | Coverage | Illumina Sequencing Platform | Accession IDs |
| NA12878 | 110x | HiSeq2500 | SRR098401 |
| HG00284 | 110x | NextSeq 2000 | ERR031490 |
| HG00448 | 110x | NextSeq 2000 | ERR032022 |
| PCa | 110x | NextSeq 2000 | SRX8023389 |
| Trio Exome Samples |  |  |  |
| NA12878 | 110x | HiSeq2500 | SRR098401 |
| NA12891 | 110x | HiSeq2500 | SRR098359 |
| NA12892 | 110x | HiSeq2500 | ERR034529 |
| Z12 | 110x | NextSeq 500 | SRR21615279 |
| Z13 | 110x | NextSeq 500 | SRR21615280 |
| Z14 | 110x | NextSeq 500 | SRR21615281 |
| Z19 | 110x | NextSeq 500 | SRR21615287 |
| Z20 | 110x | NextSeq 500 | SRR21615288 |
| Z21 | 110x | NextSeq 500 | SRR21615289 |

### Pipeline

We have used our erstwhile WES pipeline[17; Figure 1] and enhanced in-house CONsensus Variant EXome (CONVEX; https://github.com/prashbio/CONVEX) pipeline which works with four open-source variant callers in detecting variants for a comprehensive and increased sensitivity (Singh et al. 2023., communicated). In summary, FastQC (https://www.bioinformatics.babraham.ac.uk/projects/fastqc/ last accessed on September 2, 2023) was used to evaluate the quality of all samples, as we examined key characteristics of the raw reads, such as their duration, quality scores, and base distribution, to judge the data efficiency and eliminate low-quality reads. The nucleotide distribution across all reads has been obtained thanks to Phred’s optimal base quality scores, and the best quality score for GC content was set to a 50% threshold. The alignment of the reads was done using the tool Bowtie2[18] followed by the conversion of sequence alignment mapping (SAM) to binary alignment mapping (BAM) and sorting of BAM by Samtools. This analysis-ready BAM file was given to 2 of the variant callers, *i*.*e*. Freebayes (https://github.com/freebayes/freebayes last accessed on May 27, 2024) and Vt (https://github.com/atks/vt last accessed on May 27, 2024) while Bcftools and VarScan requires an additional requirement of the mpileup BAM file from Samtools as input to call variants. All the variant call sets were compared with each other using awk commands on chromosome number, chromosome position and alternate allele called by the variant callers. The consensus variants obtained were channelled through downstream analysis of validating with the ClinVar database with further genomic variant filtration as per the American College of Medical Genetics and Genomics (ACMG;[19] and Association of Molecular Pathology (AMP; [20] guidelines.

**Figure 1:**
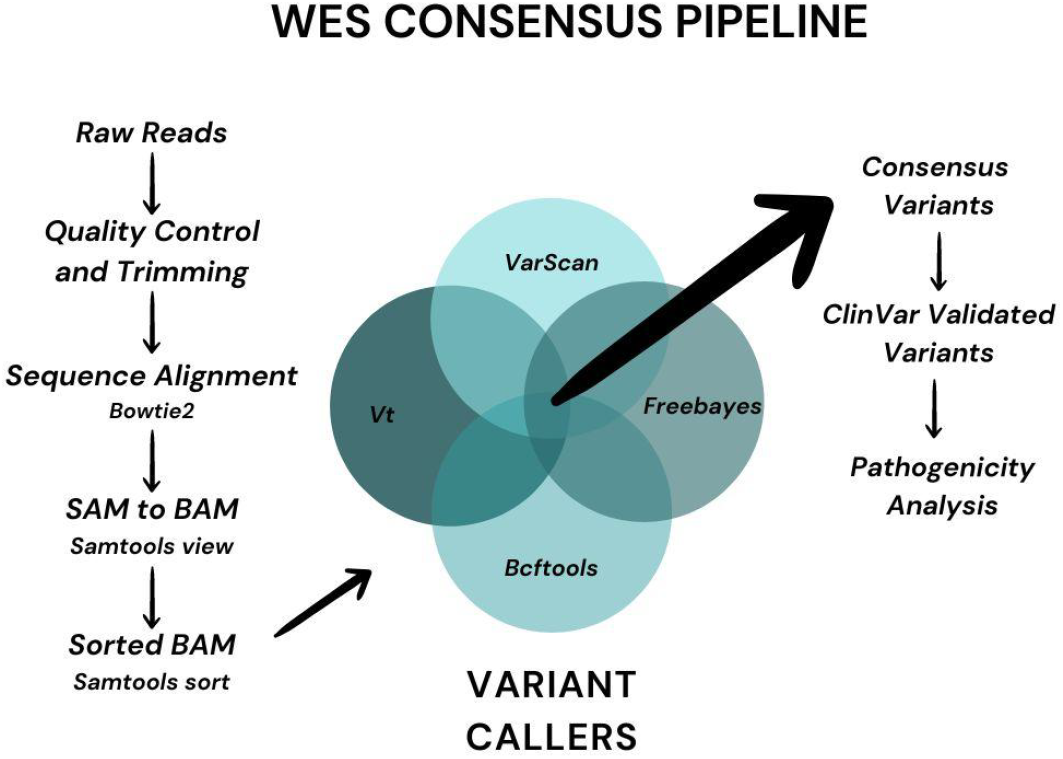
WES consensus pipeline workflow

### The Genome Analysis Tool Kit (GATK)

The GATK is a widely used tool for processing genomic data, especially for variant discovery and genotyping as it serves as the best practice for processing NGS data that are widely adopted in the genomics community[21]. After the quality control steps of FastQC and trimming of adapters with Trimmomatic[22], the read alignment was done by a high-quality aligner such as BWA-MEM to yield a SAM file[23]. The duplicates were later marked with the *Picard MarkDuplicates* to remove the duplicates that arise from PCR amplification or sequencing biases. The final step was to identify variants in the sample using GATK’s HaplotypeCaller tool[21]. Over the last few years, Unified genotyper has been deprecated in GATK even as the tool uses a probabilistic approach to assemble haplotypes and call variants simultaneously, taking into account the quality scores and alignment information for each read. Further, the raw variants obtained were filtered with various tools with the SNPs and Indels were selected through the SelectVariants tool.In this process, we considered SNPs (not Indels) for further filtering with standard filters as per the GATK best practices guidelines for the refinement of variants. The filtered SNPs were then validated with the ClinVar records and genetic variation analysis was done for benchmarking with the consensus pipeline.

### Trio-exome analysis

For the analysis of the trio-exome datasets, two variant callers having trio sample calling functionality were used (Figure 2a). The initial preprocessing of the samples was done by FastQC and Trimmomatic for all the samples in the three trio datasets. While read alignment was done using Bowtie2 for all the datasets, the individual SAM files were converted to BAM with the addition of read groups. It was further sorted and indexed with Samtools and the analysis-ready BAM files of each trio were given to the respective trio variant caller. For the VarScan pipeline, the Samtools mpileup function takes in three sorted BAM files to make one joint trio mpileup file. This file is given as input to the VarScan trio subcommand which does the variant calling with default parameters. VarScan trio utilises the family relationships and phenotypes to improve the accuracy of variant calls. It calls variants across all three samples using default parameters of --min-var-freq of 0.20 and --p-value to be 0.05. Next, it classifies any variant as Mendelian Inheritance Errors (MIE) even as it recalls such variants with adjusted settings and corrects them or the site will be marked as MIE or DENOVO. The joint variant calling workflow by GATK is a comprehensive method to detect genetic variants from sequencing data accurately. It consists of three main steps: Haplotype Caller, CombineGVCFs, and GenotypeGVCFs.The initial step, Haplotype Caller, uses local *de novo* assembly to identify variants in individual samples by reconstructing haplotypes and saving the results in the genomic Variant Call Format (gVCF). In the gVCF format, variant sites are preserved, while non-variant sites are grouped into blocks based on genotype quality during the calling process. This method allows for compression of the Variant Call Format (VCF) file without excluding any sites, thus facilitating efficient joint analysis in subsequent stages. We used the family information, such as a pedigree file and gVCF of all samples in the trio set (the father, mother, and afflicted kid), to call *de novo* variants. To identify *de novo* variants in the trio cohort, we initially generated gVCF files for each sample within the trio using the HaplotypeCaller tool. These individual gVCF files were then merged into a multi-sample gVCF file using CombineGVCFs. Subsequently, GenotypeGVCFs were employed to call the raw variants. The result of running GenotypeGVCFs is a consolidated VCF file that contains genotype information for every variant site across all the samples. This file includes details about the genotypes, qualities of genotypes, and other pertinent annotations. GenotypeGVCFs provide flexibility in handling datasets with various ploidy levels and are capable of accommodating diploid, haploid or polyploid genomes. After this, a genotype refinement workflow (Figure 2b) was implemented which included three tools obtaining the posterior probabilities of genotypes through CalculateGenotypePosteriors, and VariantFiltration, employed to remove low-quality genotypes. Additionally, VariantAnnotator annotated *de novo* SNVs and the filtered SNVs from both pipelines were analysed for pathogenicity through concordance with the ClinVar database records and our results were compared with the previous study. Finally, the benchmarked datasets on consensus and GATK pipelines were analysed with a truth set. For additional verification of the ability of the developed consensus pipeline in comparison to the established Gold Standard GATK pipeline, both pipelines were run on a widely used benchmarking dataset, NA12878 exome sequencing data. The availability of truth set for variant calls given by the GIAB acts as a reference to which both the pipelines could be compared and the consensus pipeline evaluated without any bias.

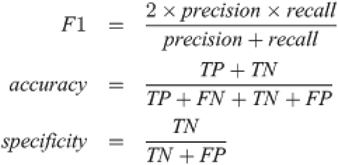

**Figure 2:**
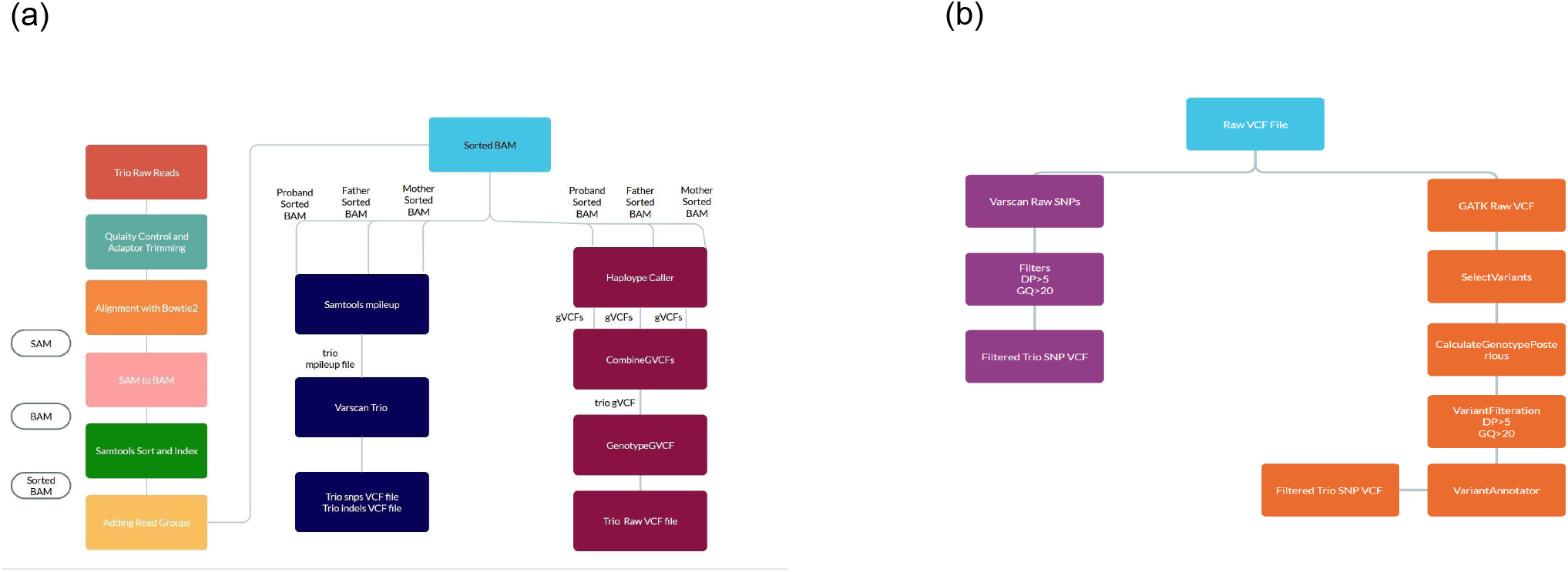
(a) An overview of steps used in the trio-exome pipeline with VarScan and GATK and (b) Genotype refinement workflow

### Statistics

We assessed the performance of our pipeline using three key metrics: F1 score, sensitivity, and specificity. The F1 score combines precision and recall, providing a balanced evaluation of our model’s accuracy. Sensitivity measures the correct identification of positive cases, while specificity measures the correct identification of negative cases. These metrics were computed for each experimental run to evaluate our method’s performance and compare it with other approaches. By utilising the F1 score, sensitivity, and specificity metrics, we aimed to evaluate our method’s performance and determine its suitability for the task at hand.

## Results and Discussion

### The WES consensus pipeline yielded a significant number of variants

The pipelines were executed on a server equipped with a 40-core CPU, 128 GB of RAM and L3 cache. The execution time for one exome sequenced sample was approximately 6 h 30 mins for GATK and 5 h 30 m for the consensus pipeline. The time required for variant calling was approximately 3 h 30 mins for GATK and 1 h 30 mins for consensus pipeline wherein on an average 3,00,000 variants were identified in each sample. We observed that the GATK called more SNVs and indels, but the consensus pipeline gave a higher average transition-to-transversion (Ti/Tv) ratio (5.57 for consensus and 3.45 for GATK). The Ti/Tv ratio is an important criterion for assessing the quality of SNV calling, for exome sequencing, the expected Ti/Tv ratio is approximately 3.011. A higher Ti/Tv ratio usually indicates a higher accuracy of SNV calling.

The genetic analysis of three samples, HG00448, HG00284, and PCa revealed significant variations in SNVs and Ti/Tv ratios (Table 2; Figure 3). The GATK pipeline consistently yielded higher SNV counts across all samples. Ti/Tv ratios indicated a higher prevalence of transitions in consensus SNVs, whereas the GATK SNVs showed a more balanced distribution between transitions and transversions. While we observed that the consensus pipeline yielded a large number of variants with conflicting pathogenicity, the variants called from both pipelines were further evaluated for pathogenicity based on the presence of variants in the ClinVar database and the concordance check of the identified variation in clinical significance (Table 3).

**Table 2:**
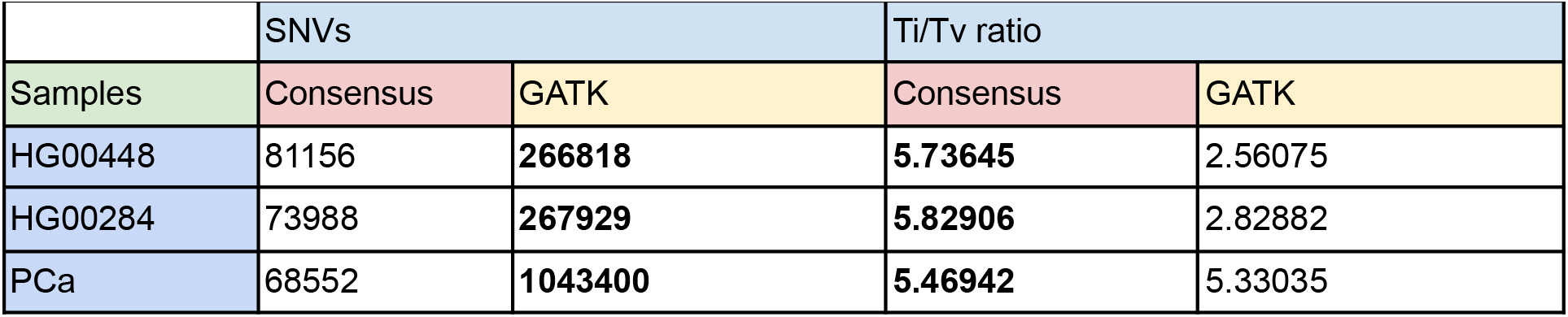
Variant counts and Ti/Tv Ratio from the two pipelines.

|  | SNVs |  | Ti/Tv ratio |  |
| --- | --- | --- | --- | --- |
| Samples | Consensus | GATK | Consensus | GATK |
| HG00448 | 81156 | **266818** | **5.73645** | 2.56075 |
| HG00284 | 73988 | **267929** | **5.82906** | 2.82882 |
| PCa | 68552 | **1043400** | **5.46942** | 5.33035 |

**Table 3:**
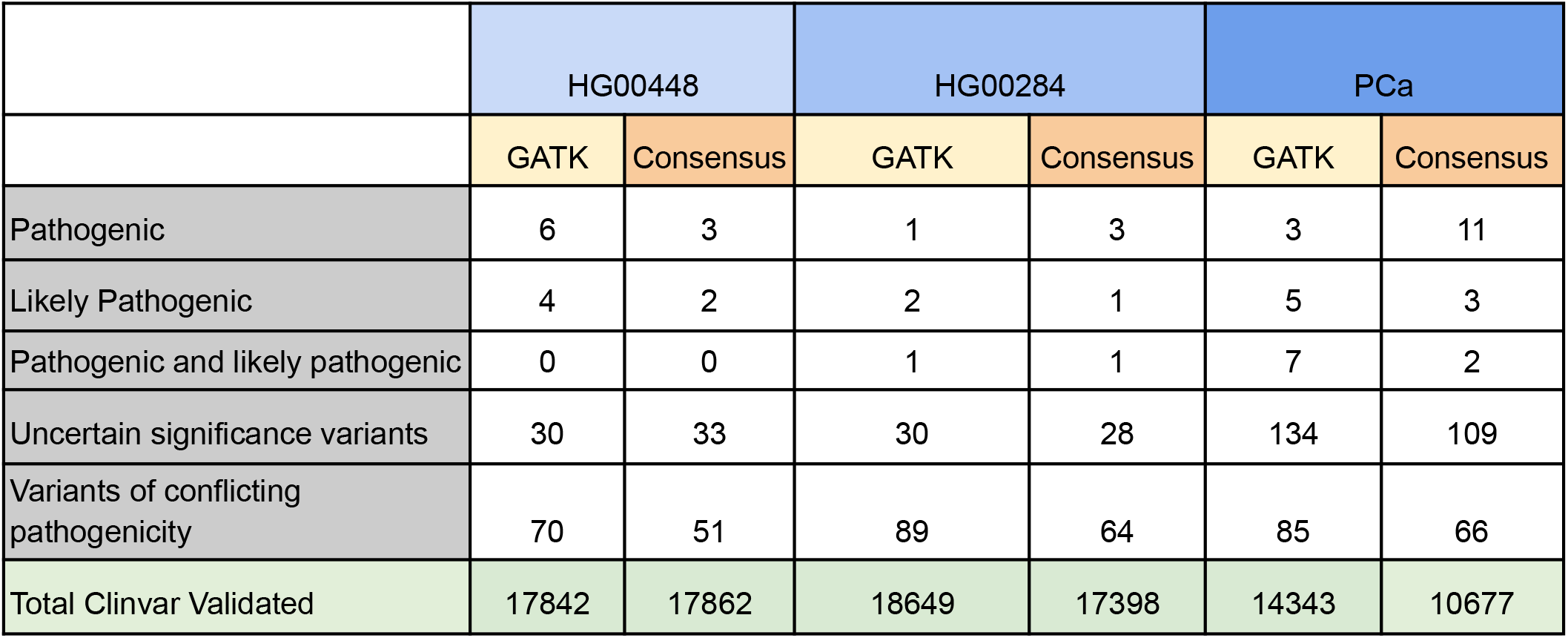
Clinical significance of variants called by the consensus and GATK pipeline.

**Figure 3:**
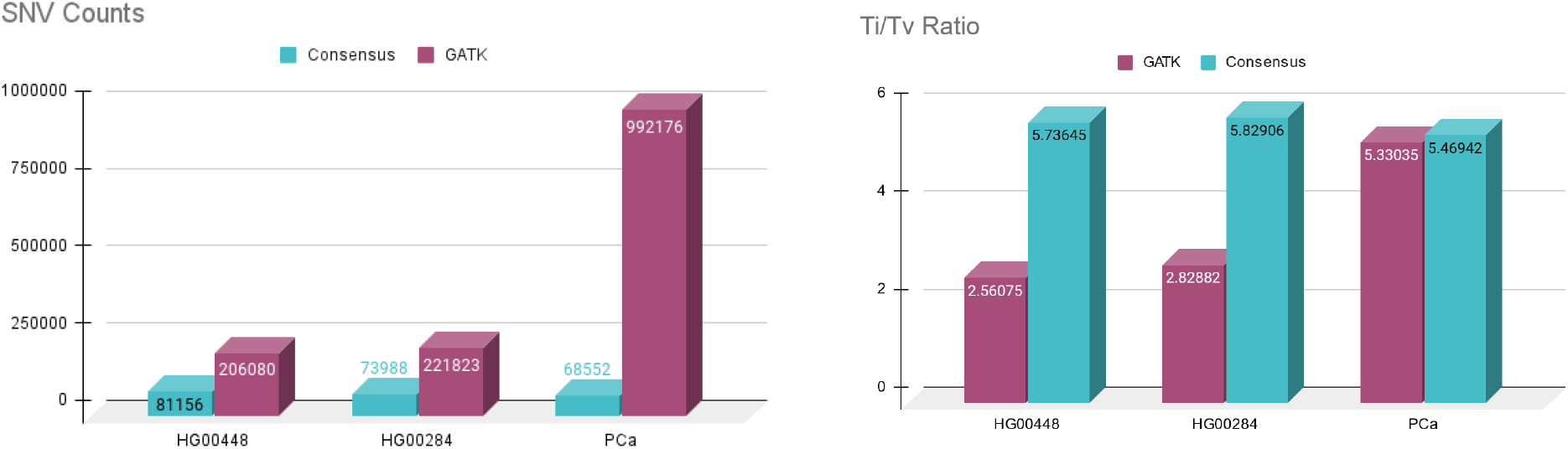
(a) SNV counts taken together from consensus and GATK pipeline and (b) Ti/Tv ratios obtained from consensus and GATK pipelines

The counts of variants for each category of clinical significance, *viz*. pathogenic, likely pathogenic, VoUS and conflicting pathogenicity was tabulated wherein our analysis showed an almost comparable number of ClinVar validated records from both pipelines. This asserts that there was a significant difference in the initial number of SNPs called by each pipeline where the final analysis resulted in a similar performance of both pipelines. We argue that the consensus pipeline appeared to be more sensitive in detecting pathogenic and likely pathogenic variants in the PCa sample, while GATK showed better performance in uncertain significance variants and ClinVar-validated variants in certain samples. The consensus pipeline also demonstrated lower numbers of variants of conflicting pathogenicity, indicating more consistent results. On the other hand, the SNPs called by GATK and the consensus pipeline were comparable and fewer in the consensus pipeline can be attributed to the default variant filtering applied by each variant caller and the algorithms used to call the variants. So the variants obtained were deemed the most confident ones among the four variant callers. The high SNP count for the truth set of NA12878 can be due to the various samples used for making the truth set with no filtrations applied.

### Benchmarking yielded variant calls from both the pipelines with ClinVar validation

Benchmarking workflows include statistical metrics such as sensitivity, precision, F1 score and specificity which provide an insight into the pipeline performance. Here both the GATK pipeline metrics and the consensus pipelines are compared against the truth set variants of benchmark NA12878 dataset. The variant calls obtained from both pipelines were further annotated with the ClinVar database for the clinical significance of the genetic variants providing clinical interpretations for variants (Table 4). For the current analysis, this annotation will help identify the various metrics associated with the benchmark workflow to assess the pipeline’s ability to call clinically relevant variants as well as concordance with the established pipelines.

**Table 4:** Pathogenicity analyses for GATK, consensus and truth set.

|  | NA12878 |  |  |
| --- | --- | --- | --- |
|  | GATK | Consensus | Truth set |
| Pathogenic | 6 | 5 | 9 |
| Uncertain significance variants | 113 | 78 | 112 |
| Variants of conflicting pathogenicity | 110 | 95 | 116 |
| Clinvar Validated | 22062 | 14809 | 34074 |

The consensus pipeline had a total of 14809 ClinVar validated variants wherein 5 of them were pathogenic, 4 were identified as true positives and 1 as a false positive. While specificity was found to be 0.999971, the sensitivity of the consensus pipeline was 0.444, and the F1 score was around 0.571. On the other hand, the GATK pipeline, which processed a larger dataset of 22062 ClinVar validated variants, identified 6 pathogenic variants. It achieved 5 true positives and 4 false positives, yielding a higher specificity of approximately 0.999882. The GATK pipeline demonstrated a higher sensitivity of around 0.833 and an F1 score of approximately 0.667.

### Comparison of trio-exome pipelines yielded distinct variants

One of the three trio-exome datasets is a 1KGP benchmark dataset that is a female and her father and mother were taken as a trio for this analysis. The other two families are CPC families(Z12 and Z19) in which only the proband is affected with the disease with the parents unaffected. The GATK trio pipeline took approximately 17 hours to finish one trio set and the VarScan trio pipeline took approximately 8 hours to complete the trio workflow. Both the pipelines included various steps of joint variant calling of trio samples and that is probably the reason for the increased time for the pipeline completion. GATK also employed additional genotype refinement as well as some joint calling specific subcommands which increases the accuracy and specificity of the variants called but also increases the total time taken to finish the pipeline.

The GATK and VarScan pipelines called distinct variants even as GATK relatively called significantly more variants than VarScan probably because of its sensitivity-focused algorithm (Table 5; Figure 4). Nonetheless, both pipelines require a more stringent filtering approach which was carried out in the subsequent steps. On the other hand, these pipelines may have a false positive rate due to potential filtering artefacts. GATK also employs local realignment and haplotype assembly techniques to handle regions with complex variation, such as indels accurately. This approach enables GATK to detect and call variants that might perhaps be missed by other callers. The variant annotation of the filtered variants was done with the ClinVar database variant record vcf file (Table 6). The annotations were determined to be present in ClinVar records by concordance check with chromosome number, chromosome position, reference and alternate allele in both the pipelines vcf files and ClinVar records. As expected, the counts for pathogenic and likely pathogenic variants were very less as two of the samples are rare diseases.

**Table 5:** Total variants called by VarScan and GATK trio pipelines.

|  | Total SNVs |  | After filtration SNVs |  |
| --- | --- | --- | --- | --- |
| Samples | GATK Trio | VarScan Trio | GATK Trio | VarScan Trio |
| NA12878_family | 1788766 | 70064 | 677150 | 49051 |
| Z12_family | 1326375 | 145238 | 277160 | 102124 |
| Z19_family | 1408848 | 152132 | 332480 | 108207 |

**Table 6:** Clinical significance of variants from Trio pipelines.

| Samples | Pathogenic | Likely Pathogenic | Uncertain significance variants | Variants of conflicting pathogenicity |
| --- | --- | --- | --- | --- |
| NA12878_family | 8 | 6 | 198 | 217 |
| Z12_family | 10 | 6 | 245 | 264 |
| Z19_family | 13 | 5 | 236 | 223 |

**Figure 4:**
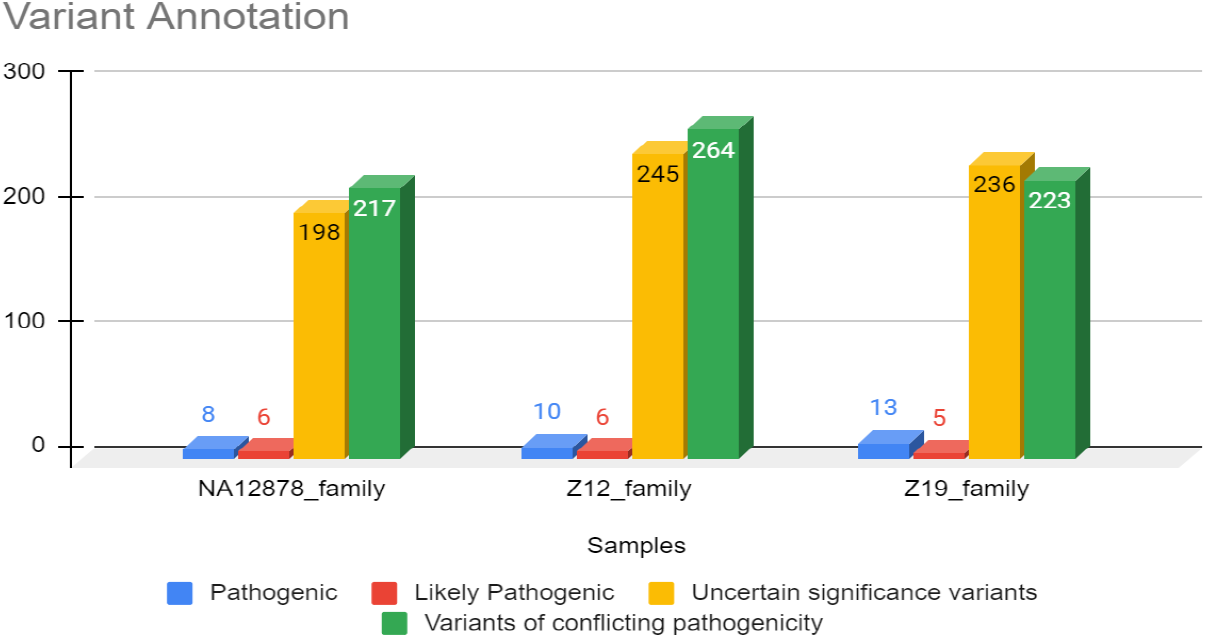
Count of variants after pathogenicity analysis for truth set analysis

### Functional annotation of selected variants

The variants were analysed using The Genome Aggregation Database (gnomAD), gave us additional information on MAF besides Combined Annotation Dependent Depletion (CADD) scores which are key factors when analysing variants in a genome, particularly in the context of rare diseases. The accepted MAF threshold for defining rare variants in the context of rare diseases is generally considered to be less than 1% and we deemed it to identify extremely rare variants (MAF<=0.01%). Nonetheless, CADD scores are generally calculated as phred-like scores and higher scores may indicate more deleterious mutations (CADD>=10; Table 7). We couldn’t consider Genomic Evolutionary Rate Profiling (GERP) scores keeping in view of the already existing stringent threshold we set.

**Table 7:**
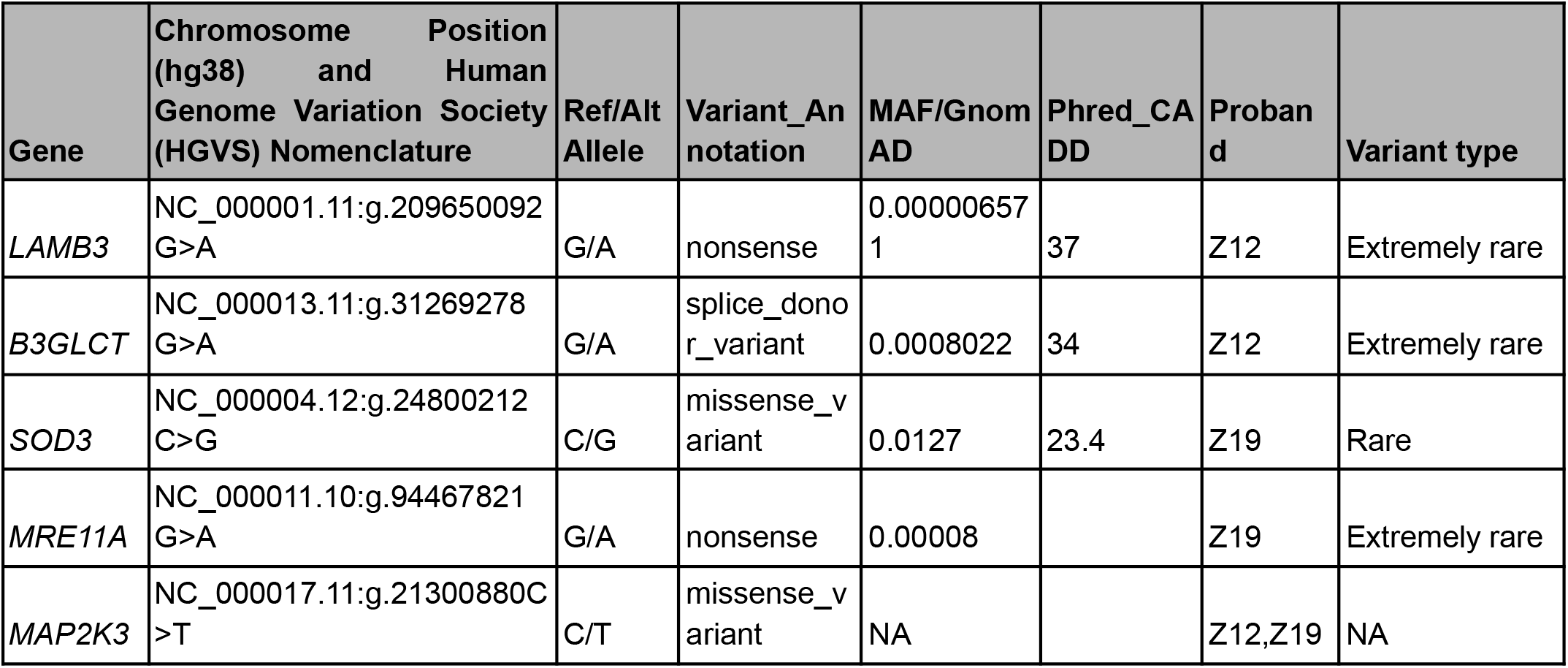
Functional annotation of selected trio variants.

Our analyses revealed the presence of novel variants involved in key cellular pathways, *viz*. cell proliferation, differentiation, migration and tumorigenesis. As anticipated, the occurrence of pathogenic and likely pathogenic variants was relatively limited, considering the rarity of conditions like CPC, a disease characterised by inflammation, fibrosis and hyperplasia, even as several genes encompassing a range of functional roles were identified. For example, *SOD3* participates in repairing oxidative damage, *MRE11A* is involved in DNA double-strand break (DSB) repair, and LAMB3 plays a role in cellular organisation and migration. *MAP2K3* regulates cell proliferation and invasion and was found to be coexpressed with other MAP kinase pathway proteins, *viz. MAP3K3*. Two SNPs in the *MAP3K3* gene were candidates for association with colon and rectal cancers while *LAMB3* is known to mediate attachment, migration and organisation of cells into tissues during embryonic development, in addition it is involved in the invasive and metastatic abilities of some types of cancer, including colon cancer. However, none of the mutations from our erstwhile exome-trio analysis which reported the significant/extremely rare variants were identified in our pipeline.

The present study attempts to benchmark the consensus exome pipeline by comparing it with a common and widely used gold standard GATK, which comes with standard best practices guidelines which help in benchmarking custom pipelines. Four datasets were taken for evaluation, three 1KGP datasets and one PCa sample. The high confidence variant call set from NA12878 was mainly used for comparing and benchmarking the consensus pipeline along with two other 1KGP samples and one PCa sample. This was performed because different variant callers may have varying strengths and weaknesses in detecting specific types of variants, such as SNVs, insertions, deletions, or structural variants. By integrating multiple variant callers in a consensus pipeline, we can improve the overall sensitivity of variant detection. Variants called by multiple callers are more likely to be true positives, reducing the risk of missing important variants. On the other hand, the consensus pipeline used four open-source variant callers at its core and all four of them contributed to the variant calls made. In the benchmarking studies, although it called fewer variants than the gold standard GATK pipeline, the Ti/Tv ratio was found to be consistently greater than what GATK achieved. This shows that with less number of calls, the consensus pipeline was able to capture a greater quality of SNVs and have a better ability to differentiate between true genetic variations and sequencing artefacts, particularly in the context of exome sequencing. On the other hand, GATK called more SNVs and indels, which could be attributed to differences in variant calling algorithms and default variant filtering applied by each pipeline. Analysis of ClinVar validated variants showed that both pipelines yielded a similar number of validated records, indicating a comparable performance in identifying clinically relevant variants. The consensus pipeline demonstrated higher sensitivity in detecting pathogenic and likely pathogenic variants in the PCa sample, while GATK performed better in uncertain significance variants. The consensus pipeline also showed lower numbers of variants of conflicting pathogenicity, suggesting more consistent results. To validate the consensus pipeline against the established gold standard GATK pipeline, both pipelines were run on a widely used benchmarking dataset (NA12878 exome sequencing data) with a truth set for variant calls indicating an unbiased evaluation and comparison of the consensus pipeline’s performance. The results showed comparable SNP calls between the pipelines, with fewer SNPs in the consensus pipeline possibly due to default variant filtering and the selection of the most confident variants. Benchmarking workflows, including sensitivity, precision, F1 score, and specificity, were used to assess the pipeline’s performance, and both pipelines were compared against the truth set variants. The clinical significance annotation with the ClinVar database further aided in evaluating the pipeline’s ability to call clinically relevant variants.

Our work, however, has inherent limitations. We could optimise the use of additional variant callers or incorporate new algorithms that specialise in detecting specific types of variants, such as indels or structural variants. Fine-tuning the variant filtering step within the consensus pipeline may also enhance its ability to differentiate true genetic variations from sequencing artefacts, thereby increasing the accuracy of variant calling. To further validate the consensus pipeline, it would be beneficial to run it on a diverse range of benchmarked datasets and compare its performance against other established gold standard pipelines. This would provide a comprehensive evaluation of its strengths and weaknesses and establish its reliability across different datasets and variant calling scenarios. Incorporating machine learning and artificial intelligence techniques into the consensus pipeline could also enhance its performance. By leveraging these approaches, the pipeline could learn from large-scale datasets and improve its ability to accurately identify and annotate clinically relevant variants. Additionally, integrating external databases and resources beyond ClinVar, such as other disease-specific databases or functional annotations, would further enrich the pipeline’s clinical significance assessment and provide more comprehensive information for variant interpretation. Nonetheless, we have a persistent dearth of trio-exome samples in the public domain supporting this limitation. Furthermore, Google’s DeepVariant[24] and Speed-seq [25] could provide deep insights as comprehensive evaluation for SNPs referred from molecular consequences are obtained. As systematic benchmarking of such variant calling pipelines is underway for variant discovery[26], we have a firm hope that there is a dawn of a new beginning in the trio-exome analysis.

## Conclusions

We aimed to evaluate the consensus exome pipeline on four datasets by comparing it to the widely used GATK, which serves as a gold standard. The consensus pipeline consistently exhibited a higher average Ti/Tv ratio, indicating improved accuracy in identifying SNV calls suggesting that the consensus pipeline excels in distinguishing genuine genetic variations from sequencing artefacts, particularly in the context of exome sequencing. It also demonstrated greater sensitivity in detecting pathogenic and likely pathogenic variants in the PCa sample, while GATK performed better with VoUS. Moreover, the consensus pipeline proved to be efficient, requiring less computational time and expertise for replication. Its comparable results to the gold standard GATK pipeline underscore its reliability and effectiveness in variant analysis. Overall, the consensus pipeline emerges as a valuable and efficient tool for variant calling, exhibiting promise in diverse sequencing data scenarios and in identifying clinical significance for relevant diseases. For the trio exome analysis, GATK has a significantly higher number of variants compared to VarScan because of its sensitivity-focused algorithm. To annotate the identified variants, we utilised the ClinVar database, which provides comprehensive data on known genetic variants and their associations with diseases. We envisage that the consensus pipeline demonstrates great potential for variant calling and clinical significance annotation, and further research and development efforts will help unlock its full capabilities in the field of genomics and precision medicine. Evaluating emerging tools and methodologies in genomics can provide valuable insights with increasing sample size and diversity of studied trios to improve understanding of rare diseases and their genetic basis. On the other hand, functional annotations and predictive models aid in prioritising pathogenic variants. For this, collaboration among researchers and clinicians, along with data-sharing initiatives and centralised databases to address the challenge of limited variants in rare diseases is needed. These approaches enhance accuracy, discover novel disease-associated variants and improve diagnosis and treatment.

## CRediT author statement

RMR and UPS contributed equally to the work. RMR benchmarked the pipelines while UPS identified the datasets. UPS tweaked our previous WES pipeline which is now called CONVEX. PS ideated, supervised and curated the project, and proofread the manuscript.

## Acknowledgements

The authors gratefully acknowledge the chancellor, revered Mata Amritanandamayi Devi and Bipin G Nair, Dean, Amrita School of Biotechnology, Amrita Vishwa Vidyapeetham for their support. The authors thank Bhargavi R. for tabulating Ti/Tv. PS sincerely thanks his collaborators, *viz*. Krishna Mohan Medicherla, Praveen Mathur, Sonal Gupta and Obul R Bandapalli for the trio-exome project that they were associated with.

## Funding

None

## Conflicts of interests

The authors declare no competing interests whatsoever. PS is a Founder of Bioclues.org, India’s bioinformatics society working for mentor-mentee relationships.

## Data availability

Our CONVEX pipeline is available at https://github.com/prashbio/CONVEX;. The source/input data accession numbers are given in Table 1. Output files are the Clinvar validated VCF files generated for both benchmarking and trio analysis.

## Declaration

Not Applicable

## Notes

https://github.com/prashbio/CONVEX

### Competing Interest Statement

The authors declare no competing interests whatsoever. PS is a Founder of Bioclues.org, Indias bioinformatics society working for mentor-mentee relationships.

### Summary of Updates

The title is changed as per suggestions of a few reviewers and the abstract slightly modified for brevity!

